# The impact of persistent bacterial bronchitis on the pulmonary microbiome of children

**DOI:** 10.1101/181982

**Authors:** Leah Cuthbertson, Vanessa Craven, Lynne Bingle, William O.C.M. Cookson, Mark L. Everard, Miriam F. Moffatt

## Abstract

Persistent bacterial bronchitis is a leading cause of chronic wet cough in young children. This study aimed to characterise the respiratory bacterial microbiota of healthy children and to assess the impact of the changes associated with the development of persistent bacterial bronchitis.

Blind, protected brushings were obtained from 20 healthy controls and 24 children with persistent bacterial bronchitis, with an additional directed sample obtained from persistent bacterial bronchitis patients. DNA was extracted, quantified using a 16S rRNA gene quantitative PCR assay prior to microbial community analysis by 16S rRNA gene sequencing.

No significant difference in bacterial diversity or community composition (R^2^ = 0.01, *P* = 0.36) was observed between paired blind and non-blind brushes, showing that blind brushings are a valid means of accessing the airway microbiota. This has important implications for collecting lower respiratory samples from healthy children. A significant decrease in bacterial diversity (*P* < 0.001) and change in community composition (R^2^ = 0.08, *P* = 0.004) was observed between controls and patients. Bacterial communities within patients with PBB were dominated by *Proteobacteria*, and indicator species analysis showed that *Haemophilus* and *Neisseria* were significantly associated with the patient group. In 15 (52.9%) cases the dominant organism by sequencing was not identified by standard routine clinical culture.

The bacteria present in the lungs of patients with persistent bacterial bronchitis were less diverse in terms of richness and evenness. The results validate the clinical diagnosis, and suggest that more attention to bacterial communities in children with chronic cough may lead to more rapid recognition of this condition with earlier treatment and reduction in disease burden.

## Introduction

Persistent or protracted bacterial bronchitis (PBB) is a leading cause of chronic wet cough lasting more than 4 weeks in young children. PBB is particularly common in pre-school children, often developing after a viral lower respiratory infection[1, 2], but may present at any age. As a result, PBB is often either misdiagnosed as asthma or the symptoms are dismissed as being due to recurrent viral infection[1–3].

Standard treatment for PBB is high dose oral antibiotics, the cough typically taking 10 - 14 days to resolve[4]. Although the aim of therapy is to provide a definitive cure, reoccurrences are frequent and if not treated successfully may lead to bronchiectasis[1–4].

Our understanding of the role of bacteria in chronic respiratory diseases is changing rapidly. Until recently it was widely believed that the healthy lung was a sterile environment[5]. A growing body of evidence, however, indicates that the healthy airways have a resident microbiota which can vary between individuals[5–7] and can alter significantly as a result of respiratory diseases such as cystic fibrosis, COPD and asthma[8].

A previous study of children with PBB using 16S rRNA gene sequencing suggested that the bacterial communities present in their lungs demonstrated similarities to those seen in children with cystic fibrosis (CF) and non-CF bronchiectasis[9]. This study provided a useful insight into the bacterial community associated with PBB, although the control subjects were undergoing bronchoscopy for clinical indications and could not be considered to be healthy[9–11].

In this present study bronchial brushings were obtained from infants and children with a diagnosis of PBB and from healthy children who were free of any respiratory symptoms or significant previous lower respiratory tract illness. This has given the opportunity for a better understanding of the microbiota of the healthy airway in childhood, as well as insight into how it is perturbed in children with PBB. In addition, the validity of characterising the lower airways microbiome using blind brushings through an endotracheal tube as opposed to more invasive bronchoscope guided sampling has been investigated.

## Methods

All study protocols were subject to ethical approval by the Local Health Research Authority (Reference: 12/YH/0230). Full details of sampling, laboratory and analytical methods are given in S1 Appendix, Supplementary Methods.

The study subjects were all 17 years of age or younger. None of these subjects had an identified significant immunodeficiency or other conditions.

Control subjects were recruited if they were undergoing an intervention requiring endotracheal intubation but were otherwise healthy without any history of acute or chronic upper or lower respiratory tracts symptoms.

Sixteen mothers of enrolled children aged ≤ 2 years had nasal and oropharyngeal (throat) swabs taken (S1 Table).

## Sample collection

Samples were obtained at the time of a diagnostic bronchoscopy for those with PBB and opportunistically from healthy subjects undergoing planned surgical procedures. Blind brushings were obtained in both groups using a protected cytology brush inserted into the airway through an ET tube. In the PBB subjects a second sample was obtained via the bronchoscope in order to compare the results from directed and blind brushings.

Once collected, brushes and swabs were immediately stored at - 80°C prior to further processing.

## DNA extractions

DNA was extracted from both swabs and brushes using the MPBio FastDNA^TM^ spin kit for soil as per the manufacturer’s instructions.

## qPCR

Prior to sequencing total bacterial burden was measured using a quantitative PCR assay targeting the V4 region of the 16S rRNA gene and using the primers 520F, 5’-AYTGGGYDTAAAGNG and 820R, 5’-TACNVGGGTATCTAATCC.

## DNA sequencing

The bacterial community within each sample was assessed using 16S rRNA gene sequencing. Dual barcoded fusion primers were used to target the previously quantified V4 region of the gene (same primers as for qPCR assay but with appropriate barcoding. See S2 Table for barcode details). Samples were sequenced on the Illumina MiSeq platform using the Illumina V2 2x250bp cycle kit. Sequences were submitted to the European nucleotide database, project number PRJEB18478.

Downstream sequencing analysis was carried out using Quantitative Insights in Microbial Ecology (QIIME) Version 1.9.0[12]. The QIIME recommended minimum threshold of 1,000 reads was applied and samples with less than 1,000 reads were removed from further analysis. All remaining samples were then rarefied to the same minimum number of reads present.

## Statistics

All statistical analyses were carried out using R Version 3.2.2[13]. Primary analysis and pre-processing was carried out in Phyloseq[13]. Non-parametric Wilcoxon sign ranked tests were used to test significant differences between means. DESeq2[14] and Indicator species analysis[15] were used to identify Operational Taxonomic Units (OTUs) significantly associated with PBB.

## Results

Twenty four children with PBB and 20 healthy controls were successfully recruited into the study (Table 1). Nasal and throat swabs were obtained from mothers for 16 of the children; 11 of the PBB children and 5 of the healthy child controls.

**Table 1.** Table of patient demographics for cases (PBB) and controls.

|  | Cases | Controls |
| --- | --- | --- |
| Number of subjects | 24 | 20 |
| Female | 14 | 12 |
| Blind brush | 24 | 21 |
| Non-blind brush | 24 | N/A |
| Age in years, mean (min, max) | 4.3 (0.8, 13.7) | 7.4 (1, 15.8) |
| Breastfed, count (min, max months) | 9 (0.07, 12) | 11 (0.5, 24) |
| Antibiotics, weeks since last dose (min, max) | 1, 25 | 4, 53 |
| Mother smokes | 2 | 4 |
| Father smokes | 5 | 10 |
| Both parents smoke | 2 | 4 |
| Mother sampled | 11 | 5 |

## 16S rRNA gene sequencing

A total of 146 samples were sequenced on the Illumina MiSeq. These included mock communities, PCR negative controls, kit controls and bronchoscopy brush controls (S2 Table). After quality control 143 samples were included for further analysis, comprising of a total of 8,833,294 reads from 1,393 distinct OTUs (61,771.29 +/- 85,954.18 [mean +/- SD] number of reads / OTU). Samples above the 1,000 read cut off recommended by QIIME were rarefied to the minimum number of reads found in the samples.

## Blind versus non-blind brush

PBB patients were sampled using both blind and non-blind brushing methods to test for potential differences in the bacterial community due to sampling method. The difference in the bacterial community of 21 paired samples were assessed using both alpha and beta diversity measurements. Samples were rarefied to 1,067 reads. No significant difference in alpha diversity was observed between the blind and non-blind brushes using richness (observed number of species, Z = 1.843, *P* = 0.068), Shannon-Weiner (bias towards rare organisms, Z = −0.017, *P* = 1), Simpsons reciprocal (bias toward more dominant organisms, Z = 0.261, *P* = 0.812) and evenness (Z = −0.052, *P* = 0.973) (S1 Fig).

Considering community composition no significant differences were observed between the different sampling methods (Adonis: R^2^=0.012, *P*=0.344). Hierarchical clustering using Bray-Curtis dissimilarity revealed that the samples clustered more closely between patients than within sampling groups (Fig 1).

**Fig 1.**
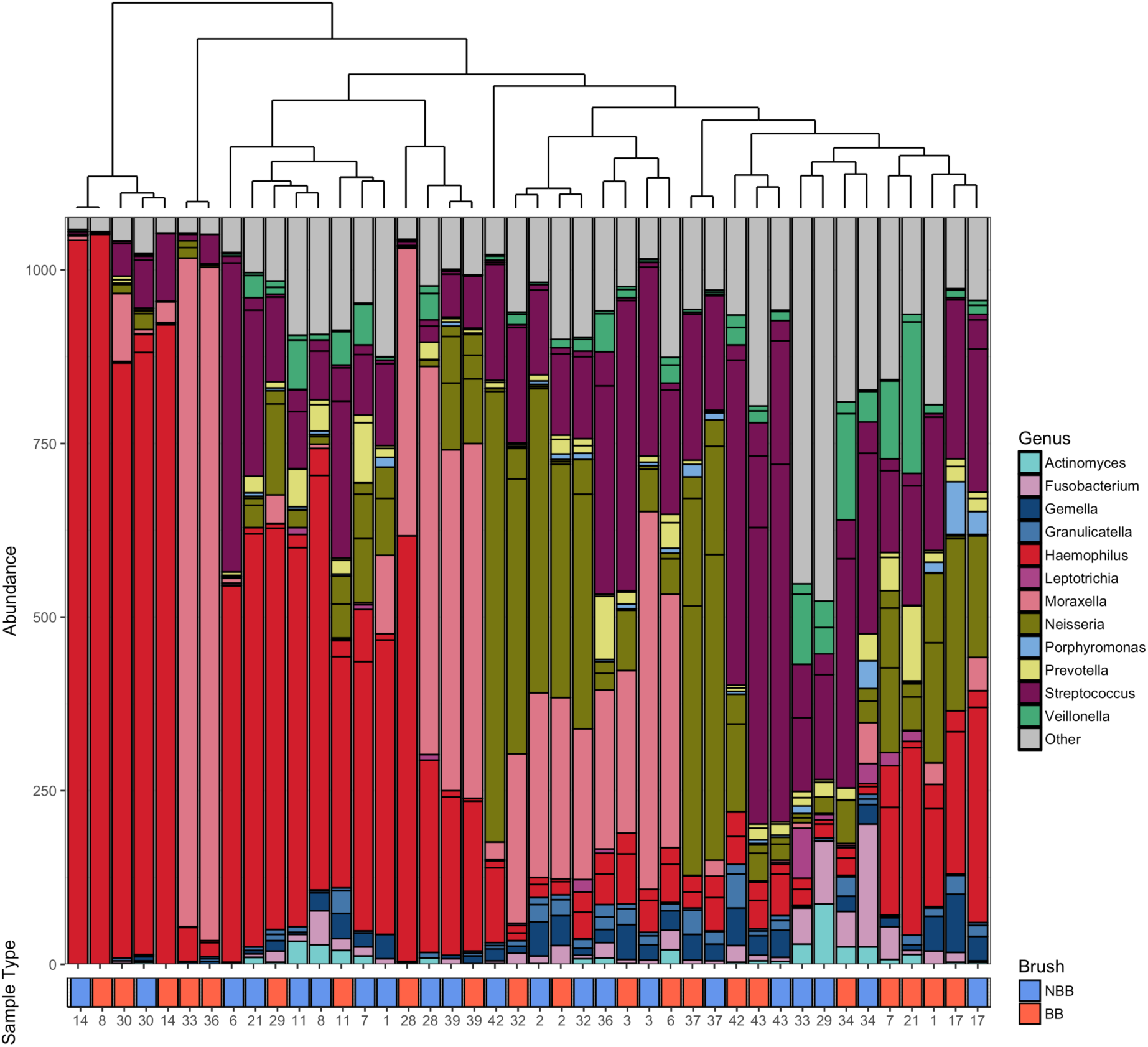
**Ordered bar chart of the top 20 OTUs present in both the blind and non-blind brushings**. Samples are ordered by a Bray Curtis dissimilarity hierarchical cluster shown by the top plot. Key to colours used for each genus is included. Identical patient numbers indicate samples were taken from the same individual. Sample type is indicated in the labelling beneath the graph with red indicating blind brush and blue indicating non-blind brush.

## PBB versus healthy controls

The bacterial community of patients diagnosed with PBB (N=24) was compared to healthy controls (N=18). Samples were rarefied to 1,150 reads, resulting in the loss of 2 control patients. No difference in the bacterial abundance by qPCR was observed between the healthy controls and PBB patients (R^2^ = 0.021, *P* = 0.511) (Fig 2). Investigation into alpha diversity measures showed that PBB patients had a significantly lower diversity than the healthy controls (S2 Fig). This was seen across all measures; the Wilcoxon rank sum test, richness (W = 100.5, *P* = 0.001), Shannon-Weiner (W = 78, *P* < 0.001), Simpson’s reciprocal (W = 79, *P* < 0.001) and evenness (W = 84, *P* < 0.001).

**Fig 2.**
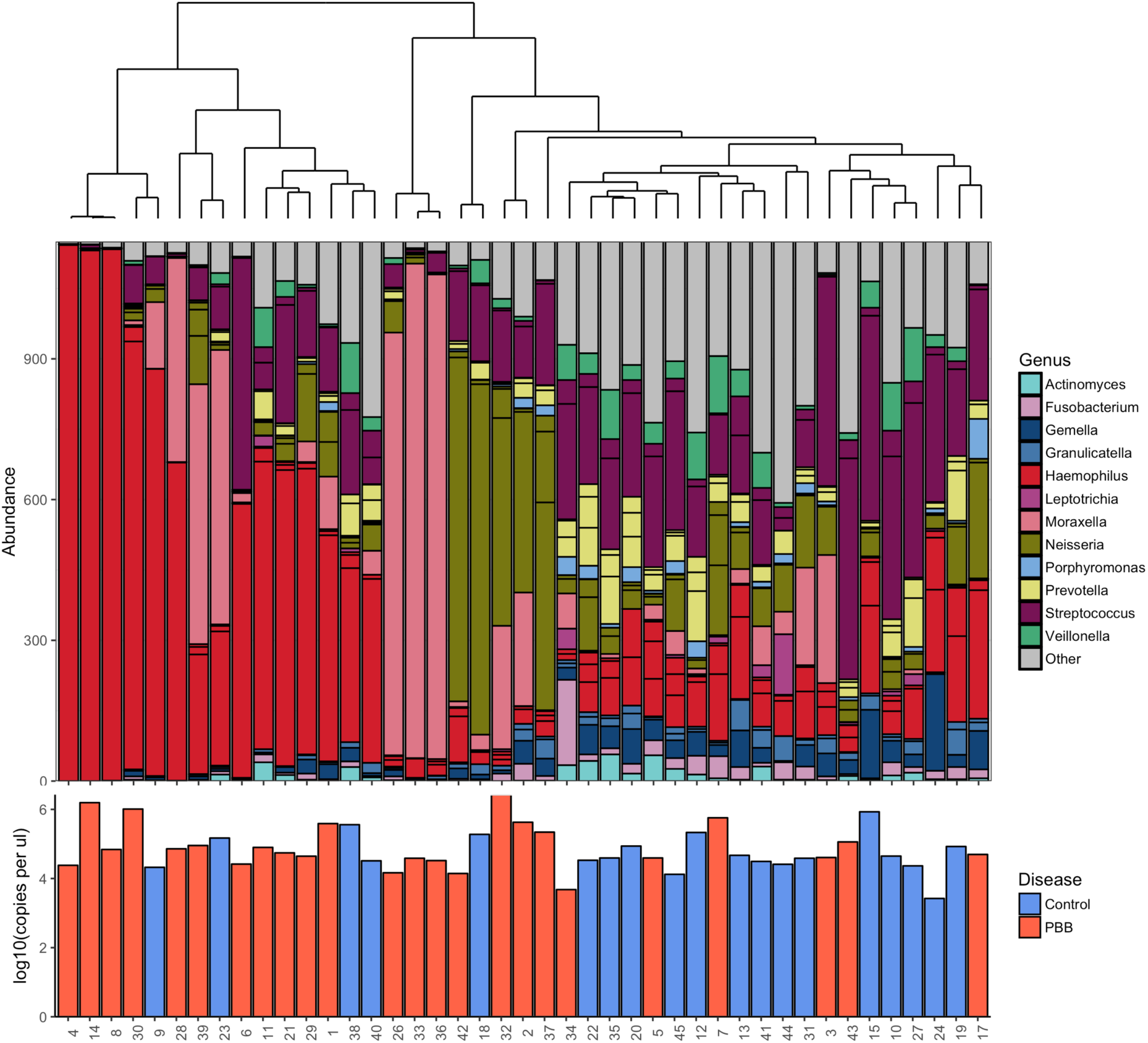
**Ordered bar chart of the top 20 OTUs present in both the PBB and control subjects.** Samples are ordered by a Bray Curtis dissimilarity hierarchical cluster, upper plot. Key to colours used for each genus is included. Patient numbers are detailed in the lower plot where disease status is also indicated by colour of the bars with red indicating PBB and blue indicates control subjects. Additionally lower bar plot indicates the log10 copies per μl as calculated by qPCR. No significant difference was found between the qPCR values between the PBB and control patients (R^2^ = 0.023, *P* = 0.445).

Differences in community composition were investigated using Bray-Curtis dissimilarity. Adonis showed significant differences in community composition (R^2^ = 0.082, *P* = 0.001). Hierarchical clustering using Bray-Curtis dissimilarity showed that the bacterial community present in healthy controls clustered separately from those with PBB (Fig 2).

DESeq2 was used to identify OTUs significantly associated with PBB. *Haemophilus* and *Neisseria* OTUs were identified as being significantly increased in patients with PBB (*P* < 0.001) (S3 Fig). This result was backed up using indicator species analysis between the PBB and control communities. Two OTUs were significantly associated with the PBB group, Haemophilus_3673 (*P* = 0.043) and Neisseria_4022 (*P* = 0.05). Haemophilus_3673 was a member of the top 20 most abundant OTUs observed. The control group had 35 OTUs significantly associated, 9 of which were included in the top 20 most abundant OTUs.

Comparing the results of standard clinical bacterial culture to the 16s rRNA gene sequencing results for the 24 PBB patients, 20 (83.33%) were culture positive. The four patients that were culture negative were, by sequencing, dominated either by *Moraxella*, *Neisseria* or *Haemophilus* OTUs. Whilst no patient cultured Neisseria, 5 of the patients were found to be dominated by a *Neisseria* OTU from the sequencing results. Seventeen (70.83%) of the 24 patients cultured *Haemophilus influenzae*, while only 9 (52.94%) of the same patients were found to be *Haemophilus* dominated by sequencing. *Moraxella* was cultured from 3 patients, in two of the three *Moraxella* was found to be the dominant organism by sequencing. Another 2 patients, however, were dominated by the *Moraxella* OTU despite not being positive on culture for the bacterium. Only fifteen of the 24 patients (62.5%) cultured the dominant organism that was identified by sequencing.

Neither parental smoking habits (R^2^ = 0.037, *P* = 0.145) or breastfeeding (R^2^ = 0.012, *P* = 0.846) were found to influence the bacterial community differences between patients with PBB and healthy controls.

To ascertain if wheeze explained any of the variation in the bacterial community observed between patients, wheeze diagnosis was investigated. No significant difference was observed between patients with and without wheeze (R^2^ = 0.048, *P* = 0.34) (S4 Fig). No control patients were diagnosed with wheeze.

PBB patients were tested for the presence of respiratory viruses. The presence of a virus (R^2^ = 0.166, *P* = 0.167) or number of different viruses present (R^2^ = 0.227, *P* = 0.095) in patients had no significant effect on the bacterial community composition. None of the 6 respiratory viruses showed any significant effect on the bacterial community composition, rhinovirus (R^2^ = 0.057, *P* = 0.266), respiratory syncytial virus (RSV) (R^2^ = 0.048, *P* = 0.406), Coronavirus (R^2^ = 0.027, *P* = 0.844), human metapneumovirus (HMP) (R^2^ = 0.052, *P* = 0.314), adenovirus (R^2^ = 0.008, *P* = 0.99), parainfluenza (R^2^ = 0.057, *P* = 0.259).

## Sampling of mothers, Throat swabs versus Nose swabs

Mothers of 16 children (11 PBB cases and 6 healthy controls) were sampled using both nose and throat swabs. To investigate if these sampling methods were comparable, samples were rarefied to 1,154 reads (4 samples were lost after rarefaction) after which non-metric multidimensional scaling (NMDS) using Bray-Curtis distance was used to investigate clustering based on bacterial community similarity (S5 Fig).

Notably samples from the same mother did not cluster together (S6 Fig). No significant difference in Shannon-Weiner (Z = −0.44, *P* = 0.7) or Simpson’s reciprocal (Z = −1.24, *P* = 0.24) diversity was observed between the two groups of samples nose versus throat swabs. There was however a significant difference in richness (Z = 2.58, *P* < 0.01) with throat swabs having more distinct OTUs than nose swabs. Additionally, bacterial abundance as determined by qPCR was significantly higher in throat swabs compared to nasal swabs (Z = 2.84, *P* < 0.01) (S6 Fig). Analysis by ADONIS confirmed these results, with 11% of the variation between samples being explained by the sample type (R^2^=0.11, *P* = 0.02).

## Children and mothers

The bacterial community within the lung of children under the age of 2 years, both with (N=11) and without a PBB diagnosis (N=5), was compared to the bacterial communities present in the nose and throat of their mothers. Samples were rarefied to 1,067 reads. Neither nose or throat swabs of mothers were found to have significantly different bacterial richness compared to child communities using a Wilcoxon rank sum test, applied to independent samples (nose; richness, W = 111, *P* = 0.317, throat; richness, W = 164.5, *P* = 0.032). Significant differences in both Shannon-Weiner and Simpson’s reciprocal were however observed between the two groups (nose; Shannon-Weiner, W = 138, *P* = 0.019, Simpson’s reciprocal, W = 138, *P* = 0.019; throat; Shannon-Weiner, W = 177, *P* = 0.007, Simpson’s reciprocal, W = 185, *P* = 0.002). Adonis showed there was significant differences in community composition when comparing between different sampling methods, while controlling for sample family (R^2^ = 0.182, *P* < 0.001), indicting that the upper respiratory tract from mothers is not comparable to their children.

Bray-Curtis hierarchical clustering was used to detect any patterns of similarities in community composition between mothers and their children (Fig 3). Overall the samples collected from the mothers compared to those from the PBB children were significantly different, however there was a single exception. The bacterial communities present in the throat swab of the mother and the blind brush from the child of family 34 were found to have in common a high abundance of S*treptococcus*, as well as *Veillonella* and *Neisseria* (Fig 6).

**Fig 3.**
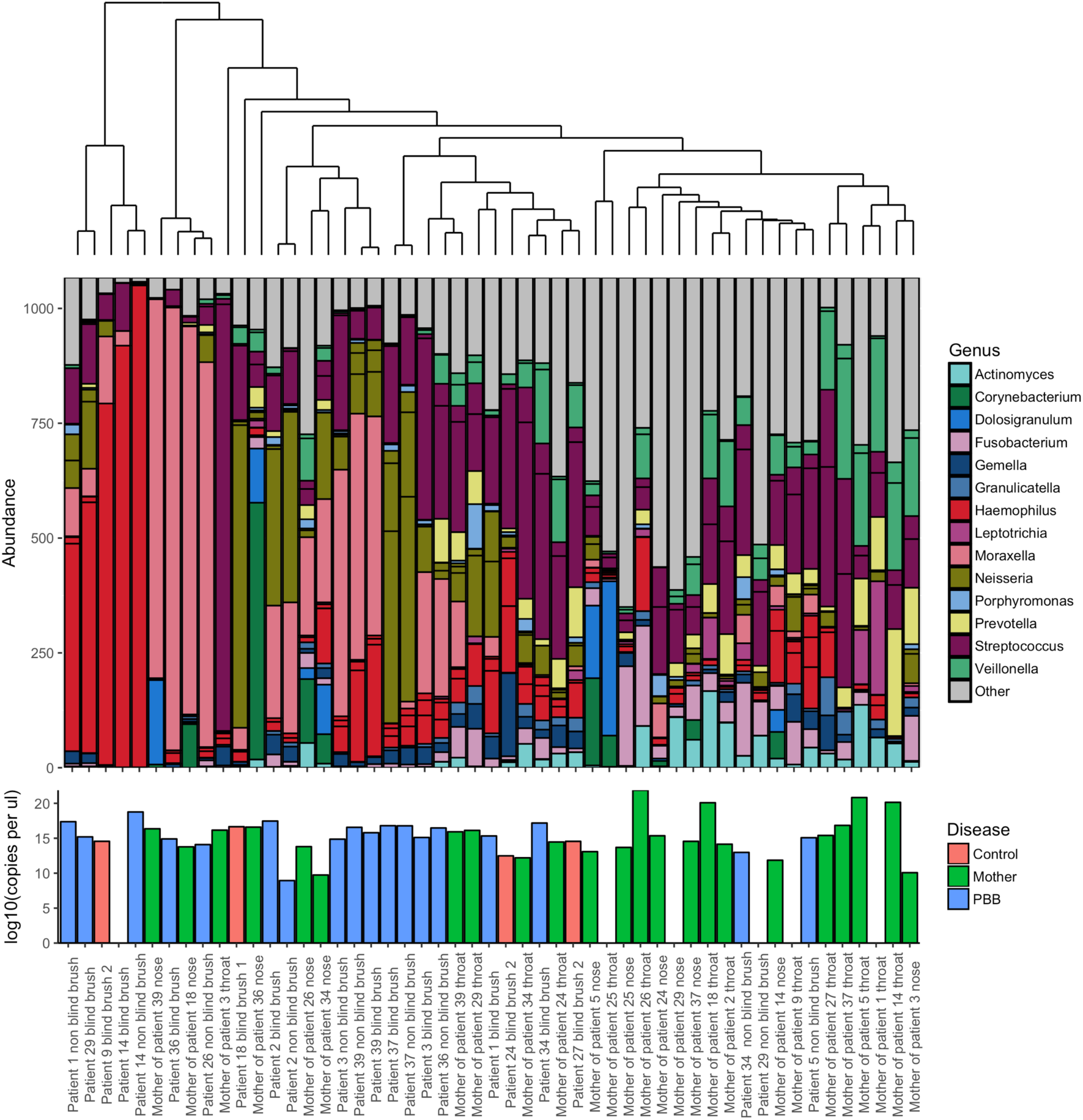
**Ordered bar chart of the top 20 OTUs present in both the mother of study children less than 2 years old and their children.** This subgroup includes both healthy children (red) and those diagnosed with PBB (blue). Mothers are indicated in green. Samples are ordered by a Bray Curtis dissimilarity hierarchical cluster, shown above. Key to colours used for each genus is included. Lower bar plot indicates the log10 copies per μl as calculated by qPCR.

## Discussion

This study provides the first insight into the composition of the bacterial community present within the lungs of both healthy infants and children and those with persistent bacterial bronchitis. Investigation into the bacterial community within the lungs of these children highlights the impact of PBB on the lung microbiota and provides insight into disease progression.

Across a range of pulmonary diseases, including COPD, non-CF bronchiectasis and cystic fibrosis, a reduction in the diversity and the appearance of dominant OTUs from potentially pathogenic genera has been observed [16–20]. It is perhaps unsurprising therefore, that in the present study a significant reduction in bacterial diversity associated with PBB was observed.

In this present study, *Haemophilus, Neisseria, Streptococcus* and *Moraxella* were all represented amongst the dominant OTUs in PBB samples, although DESeq2 and indicator species analysis showed that only a *Haemophilus* and a *Neisseria* OTU were significantly associated with PBB. This is likely to be due to the number of patients dominated in this particular sample set (Fig 3). Importantly in many cases the dominant organisms by sequencing were not those identified by culture, highlighting the potential limitations of traditional culture techniques the results of which typically dictate therapeutic strategies. It has been previously observed that conventional microbiology has the potential to miss the presence of potentially pathogenic organisms [16, 21].

Sequencing identified a Neisseria OTU as dominant in a number of subjects in this study yet culture failed to identify Neisseria in any patients. This failure to recognize organisms in chronic airways diseases may account in part for the ‘negative’ culture results from BAL and sputum samples that are frequently encountered in the presence of purulent secretions. This may be of particular importance when it comes to *Neisseria* species, which are commonly found to colonise the nasopharyngeal mucosa[22]. Despite many *Neisseria* species being considered non-pathogenic, they have been implicated in cases of pneumonia[23], periodontal disease [22] and COPD[24].

No significant differences in the bacterial community of PBB patients whose parents were smokers compared to non-smokers were found. However, due to the low number of individuals in this study we cannot exclude smoking having an effect on the bacterial communities of children. It would be important to expand this research to include a much larger sample set to investigate this further.

Wheeze is a common symptom associated with PBB [25], however in this dataset it was only diagnosed in a subset of children suffering from PBB and not in controls. No difference in the bacterial community was associated with a diagnosis of wheeze however, this may due to insufficient power.

In this study nose and throat samples were collected from mothers of children under 2 years old and compared with the results from the lower airways of their offspring. Nose swabs from mothers were significantly different from their own throat swabs, showing that there are major differences between these areas of the upper respiratory tract. Previous studies have shown that unlike the nose the bacterial community from the throat is more similar to that of the lower airways[11]. It was hypothesised that the bacterial community present in the respiratory tract of mothers and children would therefore potentially share similarities, due to their close proximity and shared environments when children are under 2 years of age. In the majority of cases however, the community composition of the mothers’ samples were significantly different from their children. Only a single case was observed where the maternal oropharyngeal community and their child’s lower airway bacterial communities were similar, and these samples were found to be dominated by the same organism.

The inclusion of lower airway samples from healthy controls provides a challenge, particularly in paediatric studies, due to the invasive nature of the sampling methods required to access the lower respiratory tract. This has resulted in many paediatric studies either sampling the upper respiratory tract [26] or including “controls” with other respiratory indications [9, 11]. Hilty *et al*. observed a decrease in the Proteobacteria and an increase in Bacteroidetes in controls compared to asthmatics, although only 3 of their 7 controls had no respiratory symptoms [11]. The current study demonstrates that the data generated from a blind brushing via an ET tube is comparable to that obtained by visualization using bronchoscopy, the so-called non-blind brush, allowing the inclusion of healthy children attending the hospital for reasons unrelated to respiratory symptoms.

This has important implications for future studies of the lower airways microbiota, particularly those involving infants and children, as a non-invasive method will allow greater number of subjects to be studied and affords the opportunity of conducting longitudinal studies with more regular sampling.

## Conclusion

In conclusion, the bacterial community within the lungs of children with PBB shows a significantly lower diversity than that observed in healthy children. This is due to the high levels of dominance of *Haemophilus*, *Neisseria*, *Streptococcus* and *Moraxella*. In many cases the dominant organisms by sequencing were not those identified by culture. This study is the first step in using next generation sequencing methods to increase our understanding of the bacterial community within the lungs of children with PBB. These methods have the potential to lead to quicker more effective treatments, reducing the risk of recurrence or disease progression.

## Ethics approval and consent to participate

All study protocols and informed consent procedures were subject to ethical approval by the NRES Committee of Yorkshire & The Humber - South Yorkshire (Reference 12/YH/0230). The study was conducted in accordance with the International Conference for Harmonisation of Good Clinical Practice and the guiding principles of the Declaration of Helsinki and the Research Governance Framework for Health and Social Care.

## Data Availability

Sequencing data generated during this study has been submitted to the European nucleotide archive (ENA), project number PRJEB18478, and is freely available. Data analysis scripts have been submitted to figshare, DOI: 10.6084/m9.figshare.4987016.

## Author contributions;

Study conception and design: VC, LB, WOCMC, MLE, and MFM; ethics, subject recruitment and sample acquisition VC and LB, performed the experiments VC and LC, acquisition, analysis and interpretation of data: LC; drafting and revision of the work as well as final approval of the submitted manuscript: all authors.

